# A Critical Rectal Temperature Defines Sex-Specific Leukocyte and Neutrophil Activation During Prolonged Passive Heat Exposure in Young Adults

**DOI:** 10.1101/2025.10.11.681783

**Authors:** Boan Wei, Yi Xu, Haojian Wang, Faming Wang

## Abstract

**Background:** Prolonged heat exposure disrupts immune homeostasis and can precipitate acute systemic inflammation. However, the core temperature threshold that triggers sex-specific leukocyte and neutrophil activation during passive heat stress remains undefined.

**Methods:** We studied 52 males and 58 females exposed to wet-bulb temperatures of 32–35 °C. Rectal temperature (*Trec*) was continuously monitored, and blood samples were collected at 0.5 °C increments up to 38.6 °C. Leukocyte and neutrophil counts were modeled using quadratic and segmented regression to identify inflection points of immune activation.

**Results:** Both leukocytes and neutrophils increased nonlinearly with rising *Trec* (*p* < 0.05). A critical *Trec* of 38.1 °C—closely aligns with the 38.0 °C occupational core temperature limits—marked an inflection in leukocyte responses: below this point, females showed steeper increases; whereas above it, males showed accelerated activation. Neutrophils demonstrated consistently greater mobilization in males across the entire temperature range (36.9–38.6 °C).

**Conclusions:** A distinct core temperature threshold (∼38.1 °C) governs immune cell activation and reveals sex-dependent response patterns. This finding provides an immunological rationale for current occupational het limits and highlights the importance of integrating sex-specific considerations into protective guidelines under extreme heat.

**Graphical abstract:** 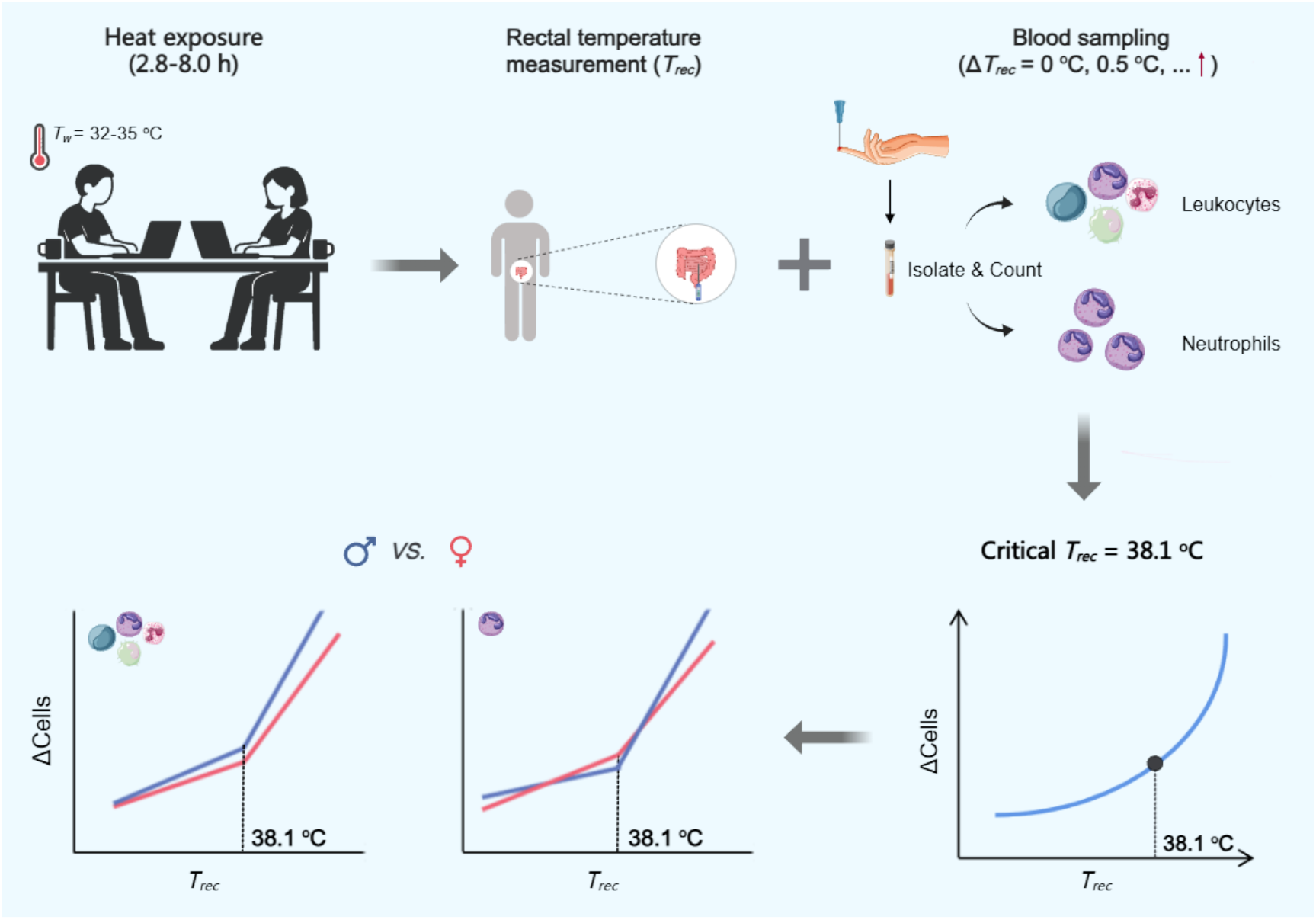

## 1. Introduction

Global warming has intensified the frequency and duration of extreme heatwaves, exposing millions to conditions that can destabilize immune and thermoregulatory homeostasis (Bouchama et al. 1993; Ebi et al. 2021; Wang et al. 2025). Even in healthy adults, sustained thermal stress can provoke systemic inflammation through activation of innate immune pathways (Bouchama et al. 2022). Leukocytes, particularly neutrophils, play a central role in initiating and amplifying inflammatory responses to thermal stress (Evans et al. 2015; Smith 1994). Yet the relationship between rising core body temperature and immune cell mobilization under prolonged passive heat exposure—without the confounding effects of exercise—remains poorly characterized.

Rectal temperature (*Trec*) is the gold-standard index of internal heat strain (Agostinelli et al. 2023; Robert et al. 2012; Towey et al. 2017). As wet-bulb temperature (*Tw*), a composite of ambient temperature and humidity, approaches critical thresholds of 33-34 °C (Schlader et al. 2020; Wang et al. 2025), evaporative heat loss is impaired, and *Trec* rises progressively even at rest (Wong 2024). Current occupational and public health guidelines designate 38.0 °C as the upper safe limit for core temperature (ACGIH, 2016; NIOSH, 2016; WHO, 1969), a value derived from cardiovascular and thermoregulatory criteria rather than immune response. Consequently, the immunological validity of this widely adopted threshold remains uncertain.

Sex-based differences in thermoregulation and immune function are well established (Klein and Flanagan 2016; Yanovich et al. 2020). Estrogen and testosterone modulate inflammatory signaling in opposing directions: females often show earlier immune activation and enhanced anti-inflammatory control, whereas males possess larger bone marrow reserves and can mount delayed but stronger responses (Bekhbat et al. 2018; Liu et al. 2022). These differences suggest that sex-specific inflection points in immune activation may emerge as core temperature rises, yet no prior work has quantified these relationships under controlled, prolonged, passive heat exposure.

Mechanistically, heat-induced leukocyte mobilization likely reflects the coordinated action of cytokines (e.g., IL-6, TNF-α, G-CSF) and chemokines that regulate both the production and directional recruitment of immune cells. G-CSF functions as a hematopoietic cytokine that stimulates neutrophil proliferation and release from bone marrow reserves, whereas classical chemokines such as IL-8 (CXCL8) and CXCL1 orchestrate leukocyte chemotaxis toward peripheral tissues (Adam and Lloyd1997; Peake et al. 2008). In parallel, sympathetic activation and hypothalamic–pituitary–adrenal (HPA) signaling modulate immune cell redistribution and priming (Dhabhar et al. 2012). Identifying the core temperature at which these cytokine–chemokine–neuroendocrine interactions accelerate provides a physiological rationale for refining current occupational heat exposure limits.

Here, we systematically exposed healthy young adults to *Tw* = 32–35 °C under 12 combinations of temperature and humidity, continuously monitoring *Trec* and sampling blood at 0.5 °C increments up to 38.6 °C. We hypothesized that leukocyte and neutrophil counts would increase in a nonlinear, temperature-dependent manner, with sex-specific differences in threshold and rate of activation. By mapping immune cell dynamics across the full thermal range, this study aims to identify the core temperature inflection point that marks the onset of systemic immune activation and to evaluate its alignment with existing occupational heat exposure limits.

## 2. Methods

### 2.1 Ethical Approval

Ethical approval was secured from the Institutional Review Board (IRB) of Xi’an University of Science and Technology (approval numbers: XUST-IRB223011 and XUST-IRB224002). All participants provided both verbal and written informed consent before enrollment, in accordance with the Declaration of Helsinki.

### 2.2 Participants and Study Design

A total of 110 healthy young adults (52 males: 24.50 ± 2.40 yr, Body mass index (BMI), 22.14 ± 1.74 kg/m^2^; 58 females: 23.20 ± 1.97 yr, BMI, 20.79 ± 2.11 kg/m^2^) participated in the study. Detailed anthropometric characteristics are presented in Supplementary Table S1. All participants underwent medical screening at a certified hospital to exclude cardiovascular, endocrine, or immune disorders, and none had engaged in occupational heat exposure or heat acclimatization training during the previous six months. Female participants were tested during the follicular phase of their menstrual cycle to minimize hormonal variability.

Participants followed a standardized pre-trial regimen for 24 hours before testing, abstaining from caffeine, alcohol, and spicy foods. On the morning of testing, baseline rectal temperature (*Trec*) wasmeasured at 08:30 h using a rectal thermistor probe (YSI401, Yellow Spring Instrument, Yellow Springs, OH, USA, accuracy: ±0.1 °C). Only individuals with *Trec* ≤ 37.1 °C were eligible to proceed; those exceeding this threshold consumed controlled volumes of iced water to normalize *Trec* before exposure. If rectal temperature remained unchanged within 30 minutes after ingestion, participants were permitted to enter the chamber, ensuring thermal stabilization before the trial began.

Experiments were conducted in a climate chamber under four wet-bulb temperature (*Tw*) conditions: 32 °C, 33 °C, 34 °C, and 35 °C. Each *Tw* level was achieved using three combinations of dry-bulb temperature (*T*db) and relative humidity, producing 12 distinct but physiologically equivalent *Tw* conditions (Supplementary Table S1). Because these combinations at each *Tw* level elicited comparable core temperature trajectories across trials (Wang et al. 2025), data were subsequently pooled across *Tw* conditions for regression modeling, ensuring adequate statistical power to detect temperature-dependent immune responses while minimizing environmental redundancy.

### 2.3 Exposure Protocol

Participants wore standardized clothing (short-sleeve cotton shirts and shorts; approximately 0.3 clo) and entered the chamber in a seated resting state. Hydration was maintained throughout exposure using a standardized protocol (8–10 mL^·^kg^-1·^h^-1^), and nude body-mass loss was maintained below 1.0% to prevent dehydration-induced hematological alterations (Supplementary Table S2). For *Tw* = 32 °C and 33 °C, exposures lasted 8 h (09:00–17:00). For *Tw* = 34 °C and 35 °C, exposure was terminated once rectal temperature reached 38.6 °C for safety. Rectal temperature (*Trec*) was continuously monitored with the YSI401 rectal thermistor probe.

### 2.4 Rationale for Capillary Blood and Sampling Protocol

Studies have demonstrated a high agreement (r > 0.97) between fingertip capillary and venous leukocyte and neutrophil counts, supporting the use of this method for repeated sampling during heat exposure (DiPasquale et al. 2025; Rao et al. 2011). As multiple blood samples (4–5 within 2.8–8.0 hours) were required, capillary sampling was chosen for its minimally invasive nature, lower participant burden, and feasibility for repeated collections without venous catheterization.

Participants remained seated prior to and during the entire exposure. Baseline sampling was performed after at least 30 minutes of seated rest upon arrival at the laboratory, and all subsequent fingertip blood collections were conducted with the participant’s arm supported on a worktable. This standardized posture minimized posture-related shifts in peripheral blood volume and ensured consistency across repeated capillary samplings. Capillary blood sampling was collected each time *Trec* increased by 0.5 °C from baseline (Δ*Trec* = 0 °C, 0.5 °C, 1.0 °C, 1.5 °C, etc.). Before sampling, the fingertip was disinfected with 75% medical alcohol and allowed to air-dry completely. A single-use 28G lancet (Sinocare, Changsha, China) was used to puncture the lateral fingertip pad; the first drop of blood, after gentle squeezing, was discarded to avoid contamination by tissue or interstitial fluids and to minimize hemolysis. Approximately 60 µL of capillary blood was collected into a microtube, divided into three aliquots (∼20 µL each), and immediately mixed with EDTA-K2 anticoagulant.

Samples were analyzed on-site within 5 min of collection using an automated hematology analyzer validated for small-volume capillary specimens (BHA-3000, Getein, Nanjing, China). The mean of three technical replicates at each *Trec* point was used for analysis. All sampling followed aseptic procedures, and the analyzer underwent routine internal quality control before and during data collection.

## 2.5 Data Analysis

Analyses were performed in SPSS 27.0 (IBM Corp., USA) and OriginPro 2025b (OriginLab Corp., USA). Anthropometric and environmental variables were summarized as mean ± standard deviation (SD). Baseline leukocyte and neutrophil counts were compared between sexes. Data normality was assessed using the Shapiro-Wilk test; as hematological variables were non-normally distributed, values were expressed as median [interquartile range (IQR): lower quartile, upper quartile]. Homogeneity of variance was tested using Levene test, and intergroup comparisons were performed using the Mann– Whitney U test. Effect sizes (*r*) were calculated to quantify the magnitude of sex differences, providing complementary information to *p*-values and reflecting both statistical and practical significance. Interpretation followed Cohen’s conventions: *r* < 0.1, trivial; 0.1-0.3, small; 0.3-0.5, medium; and > 0.5, large. Statistical significance was set at *p* < 0.05.

To characterize temperature–response patterns, increments in leukocyte and neutrophil counts were computed at each rectal temperature (*Trec*) measurement relative to baseline (Δ count = value at a given*Trec−*baseline value). The corresponding change in rectal temperature (Δ*Trec*) was calculated in parallel. These paired Δ count and Δ*Trec* data from all participants were pooled into a single dataset. Quadratic polynomial regression was then used to model the relationship between Δ counts and Δ*Trec*, yielding estimates for both the rate of change (first-order coefficient) and the acceleration of change (second-order coefficient). Model parameters were compared between sexes.

To explore potential threshold effects, segmented linear regression was subsequently applied using the critical *Trec* value of 38.1 °C identified from the polynomial models (Supplementary Text S1). This approach generated separate slope estimates for data points below and above 38.1 °C, enabling direct comparison of immune activation rates across lower and higher rectal temperature phases.

## 3. Results

Baseline leukocyte counts did not differ significantly between sexes (median: males = 6.76 [IQR 5.76– 8.02] × 10^9^ cells^·^L^-1^; females = 6.72 [IQR 5.64–7.86] × 10^9^ cells^·^L^-1^; *p* = 0.269; *r* = 0.105), indicating negligible practical differences (Supplementary Table S3). In contrast, baseline neutrophil counts were modestly higher in males than females (median = 3.13 vs. 2.99 × 10^9^ cells^·^L^-1^; *p* = 0.041; *r* = 0.199), although the effect size was small.

During prolonged passive heat exposure (FIGURE 1), leukocyte and neutrophil counts increased progressively with rising rectal temperature (*Trec*) in both sexes. From baseline to 38.6 °C, leukocyte counts increased by up to 117.2% in males and 102.4% in females, whereas neutrophils increased by 177.6% in males and 181.9% in females.

**FIGURE 1.**
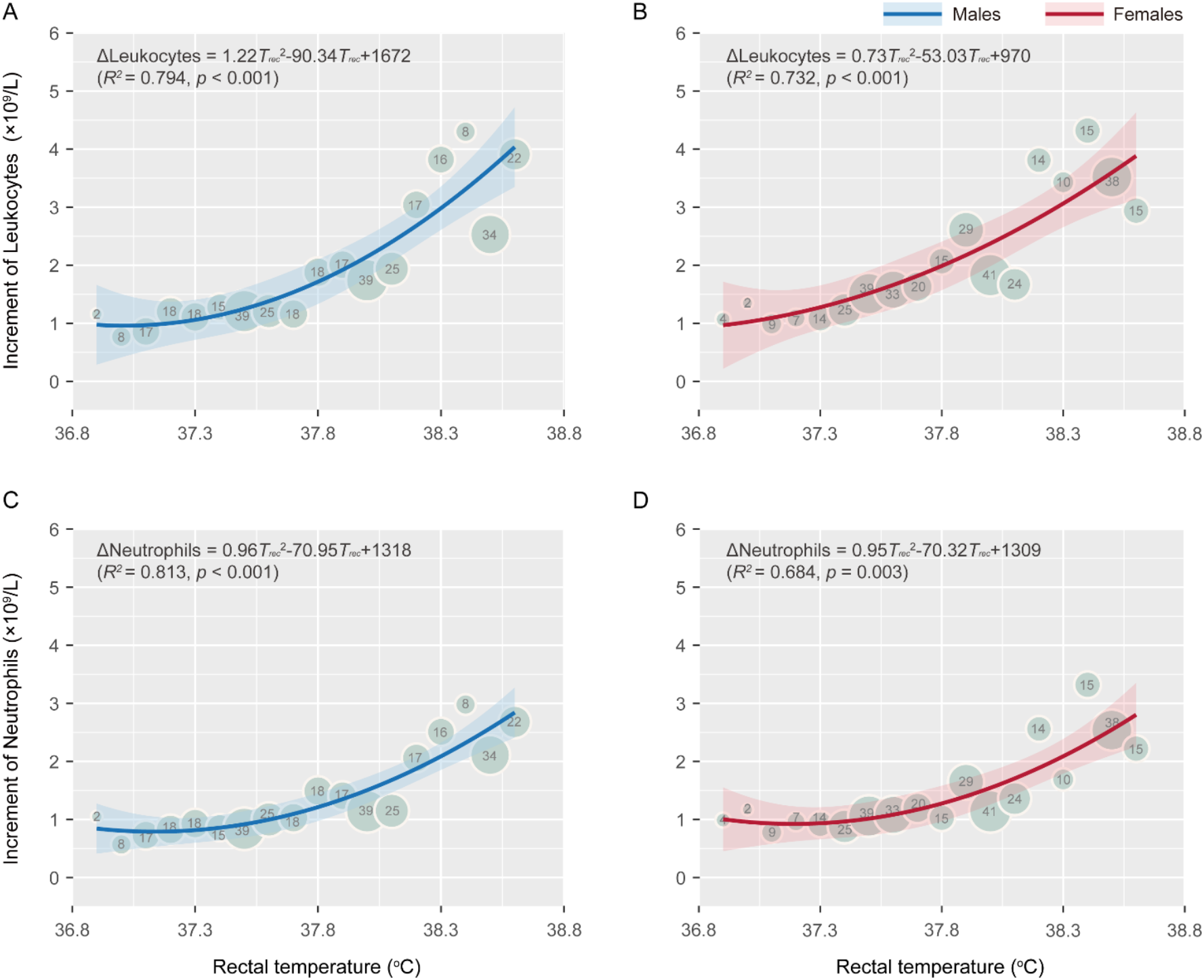
Temperature-dependent leukocyte and neutrophil changes during passive heat exposure in young adults. Quadratic polynomial regression models demonstrate nonlinear, accelerating increases in immune cell counts as rectal temperature rises from baseline (i.e.,36.9 °C) to 38.6 °C. (A) Leukocyte responses in males (n= 356 Δ leukocyte values) and (B) females (n= 354 Δ leukocyte values). (C) Neutrophil responses in males (n= 356 Δ neutrophil values) and (D) females (n= 354 Δ neutrophil values) exhibit similar nonlinear patterns, with consistently steeper slopes in males across the temperature range. Δ values represent changes from baseline (×10^9^ cells/L). Shaded regions indicate 95% confidence intervals (CIs) for fitted lines. Regression coefficient, R^2^ values, and *p*-values are shown in each panel.

Quadratic polynomial models provided strong fits for leukocyte responses in males (*R*^*2*^= 0.79, *p* < 0.001; FIGURE 1A) and females (*R*^*2*^= 0.73, *p* < 0.001; FIGURE 1B), as well as for neutrophil responses in males (*R*^*2*^= 0.81, *p* < 0.001; FIGURE 1C) and females (*R*^*2*^= 0.68, *p* = 0.003; FIGURE 1D). For leukocytes, the quadratic coefficient was larger in males (1.22) than in females (0.73), while the linear coefficient was more negative (−90.34 vs. −53.03). This pattern indicates a slower initial slope in males at lower *Trec* but a more rapid acceleration at higher temperatures. The modelled curves intersected at a critical *Trec* of 38.1 °C, beyond which male leukocyte counts increased more rapidly than female counts, consistent with the segmented regression analysis shown in FIGURE 2.

**FIGURE 2.**
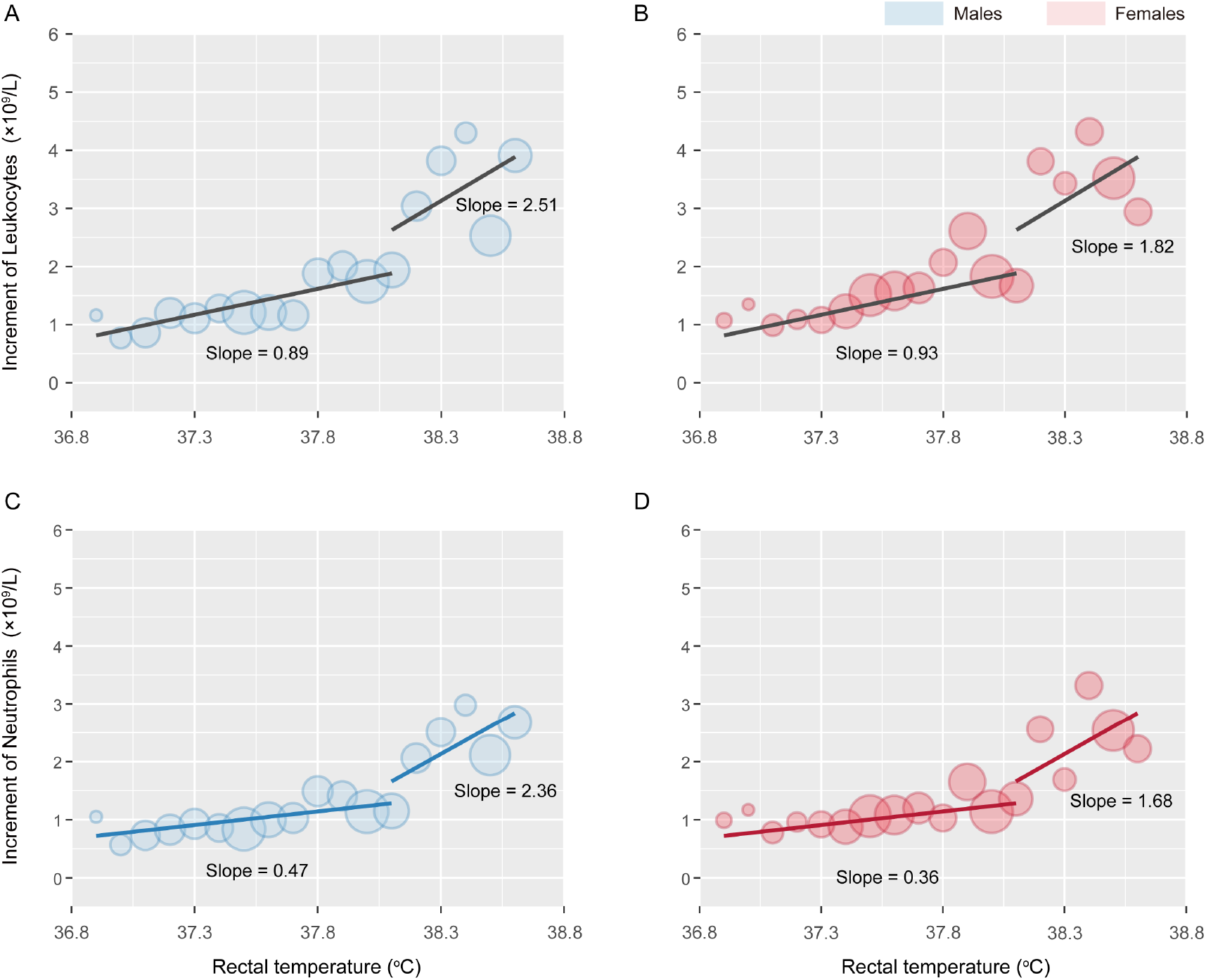
Segmented linear regression of leukocyte and neutrophil increments below and above the critical rectal temperature threshold (38.1 °C). (A) In males (n= 356 Δ leukocyte values), leukocyte slope increases from 0.89 × 10^9^ cells^·^L^-1·^°C^-1^ (below threshold) to 2.51 × 10^9^ cells^·^L^-1·^°C^-1^ (above threshold). (B) In females (n= 354 Δ leukocyte values), corresponding slopes are 0.93 and 1.82.(C) In males (n= 356 Δ neutrophil values), the slope increased from 0.47 × 10^9^ cells^·^L^-1·^°C^-1^ to 2.36 × 10^9^ cells^·^L^-1·^°C^-1^ across the threshold, while (D) in females (n= 354 Δ neutrophil values), the slope increased from 0.36 × 10^9^ cells^·^L^-1·^°C^-1^ to 1.68 × 10^9^ cells^·^L^-1·^°C^-1^. Δ values represent changes from baseline rectal temperature (i.e., 36.9 °C).

Segmented regression confirmed the threshold-dependent divergence in leukocyte responses (FIGURE 2). Below the critical *Trec* limit of 38.1 °C, leukocyte slopes were nearly identical in males (0.89 × 10^9^ cells^·^L^-1·^°C^-1^; FIGURE 2A) and females (0.93 × 10^9^ cells^·^L^-1·^°C^-1^; FIGURE 2B). Above this threshold, the male leukocyte slope increased from 2.51 × 10^9^ cells^·^L^-1·^°C^-1^ compared with 1.82× 10^9^ cells^·^L^-1·^°C^-1^ in females, a 38% greater rate of change in males. In contrast, neutrophil responses did not exhibit a curve intersection: males maintained higher slopes across the entire temperature range. Segmented analysis revealed male neutrophil slopes increased from 0.47 × 10^9^ cells^·^L^-1·^°C^-1^ to 2.36× 10^9^ cells^·^L^-1·^°C^-1^ across the threshold (FIGURE 2C), whereas female slopes increased from 0.36 × 10^9^ cells^·^L^-1·^°C^-1^ to 1.68 × 10^9^ cells^·^L^-1·^°C^-1^ (FIGURE 2D), reflecting a 40.5% higher rate in males above 38.1 °C. These results indicate both greater baseline mobilization and a steeper thermal acceleration of neutrophil responses in males.

## 4. Discussion

Understanding how core temperature modulates immune cell dynamics is critical in the context of increasing occupational and environmental heat stress. This study provides new insights into the relationship between incremental core temperature elevations and immune activation during prolonged passive heat exposure at wet-bulb temperatures (*Tw*) of 32–35 °C. Continuous rectal temperature monitoring combined with repeated blood sampling revealed nonlinear, temperature-dependent increases in circulating leukocytes and neutrophils, with clear sex differences in both activation threshold and rate. These findings extend current understanding of human physiological responses to heat stress by identifying an immunological inflection point that coincides with established thermoregulatory limits.

### 4.1 Thermal Thresholds and Immune Activation Dynamics

A critical rectal temperature (*Trec*) of approximately 38.1 °C marked a divergence in leukocyte response patterns between sexes. Below this point, females exhibited a steeper increase in leukocyte counts; above it, males showed an accelerated response with a 38.0% greater slope. This inflection aligns closely with the widely adopted occupational core temperature limit of 38.0 °C, which was originally derived from cardiovascular and thermoregulatory endpoints (Bernard et al. 2023). Our findings suggest that this physiological boundary may also correspond to a distinct shift in immune activation dynamics, adding an immunological dimension to the rationale for this limit. Whether this immune inflection precedes or parallels other systemic responses such as cardiovascular drift or perceived heat strain warrants further study, but its temporal proximity to this established threshold is notable.

### 4.2 Sex Dimorphism in Thermal Immune Responses

These observed sex differences likely reflect integrated effects of hormonal modulation, thermoregulatory control, and hematopoietic capacity. Testosterone has been shown to promote pro-inflammatory signalling via androgen receptor-mediated pathways (Gonzales et al. 2009), whereas estrogen suppresses NF-κB activation and enhances anti-inflammatory cytokine production (Villa et al. 2015). These opposing endocrine effects could contribute to the earlier leukocyte mobilization seen in females, followed by a plateau as anti-inflammatory feedback mechanisms engage.

In addition, males possess a larger leukocyte reserve pool in bone marrow (Rosenblum et al. 1974), potentially allowing for delayed but more pronounced mobilization once a critical thermal threshold is surpassed. This may explain why leukocyte activation in males accelerated more rapidly at higher core temperatures. While hormonal pathways are likely major contributors, other factors including differences in body composition, heat storage capacity, and cardiovascular strain, may also shape these sex-specific response patterns.

### 4.3 Mechanisms Underlying Rapid Leukocyte Mobilization

The rapid, temperature-dependent rise in circulating leukocytes and neutrophils can be mediated through multiple pathways that do not require concurrent cytokine elevations. Acute stressors elicit leukocyte demargination through catecholamine- and glucocorticoid-dependent mechanisms, leading to the detachment of marginated cells from the endothelium and their entry into circulation (Lemberger et al. 1996). These processes can occur within minutes and are facilitated by changes in cell deformability and vascular shear forces. On a slightly longer timescale, bone marrow release may further augment circulating cell numbers. Importantly, these mechanisms precede or occur independently of slower cytokine-driven inflammatory cascades.

This distinction has both mechanistic and applied relevance. Rapid leukocyte mobilization may represent an adaptive, transient immune redistribution aimed at maintaining host defense during acute heat stress. However, sustained activation at elevated core temperatures could amplify systemic inflammation, with potential implications for heat-related illness risk. Clinically, early immune activation may lower the threshold for inflammatory sequelae such as endothelial dysfunction, coagulopathy, or systemic inflammatory response, thereby increasing vulnerability during prolonged heat exposure.

### 4.4 Integrating Molecular Mediators and Thermoregulatory Control

Although molecular mediators were not measured in this study, previous studies have documented rises in IL-6, TNF-α, and G-CSF with increasing thermal strain (Tsai et al., 2013). Central thermoregulatory circuits and the hypothalamic-pituitary-adrenal (HPA) axis likely modulate these responses through sympathetic activation and glucocorticoid release (Douglass et al., 2023; Torres et al., 2005). Moreover, transient receptor potential (TRPV1) channels, which detect both peripheral and core heating, may serve as upstream molecular sensors linking thermal inputs to immune pathways (Lv et al., 2012). Future studies incorporating cytokine profiling, cell surface activation markers (e.g., CD62L shedding), and neuroendocrine measures would help delineate the relative contribution of demargination versus de novo activation.

### 4.5 Practical and Translational Implications

The distinct leukocyte and neutrophil activation patterns observed here underscore the importance of accounting for sex in both experimental design and heat exposure guidelines. In females, earlier leukocyte activation suggests that protective interventions—such as rest breaks or enhanced cooling— may need to occur before the conventional 38.0 °C core temperature limit is reached. In males, the rapid escalation of immune activation above this threshold raises concerns about amplified inflammatory stress during sustained exposure, even when hydration is maintained.

These findings do not imply immediate clinical risk but highlight an underappreciated physiological response domain that may interact with cardiovascular, thermoregulatory, and perceptual factors during heat exposure. Whether these immune shifts translate to increased vulnerability to heat-related illness or impaired recovery remains to be determined.

### 4.6 Limitations and Future Directions

Several limitations should be acknowledged. First, the absence of cytokine and hormonal measurements limits the ability to directly confirm mechanistic pathways underlying the observed leukocytosis. Second, the study focused on healthy young adults, and results may not generalize to older individuals or those with comorbidities, who may exhibit different thermal and immune response profiles. Third, only passive exposure was tested; exercise may potentiate these responses through combined metabolic and thermal stress.

Future research should integrate immune, endocrine, and thermoregulatory markers in real time to map the temporal sequence of these events more precisely. Longitudinal studies are also needed to evaluate whether repeated or chronic heat exposure leads to immune adaptation or maladaptation.

## 5. Conclusions

Collectively, we identified a clear, temperature-dependent inflection in leukocyte and neutrophil dynamics during prolonged heat exposure that coincides with the commonly cited 38 °C core temperature threshold. This immunological activation occurs rapidly, exhibits distinct sex-dependent patterns, and may represent an early biological signal of systemic strain. These results extend the physiological basis for occupational heat thresholds and highlight the need to consider immune responses alongside thermoregulatory and cardiovascular markers when defining safe exposure limits.

## Acknowledgments

This research was financially supported by the National Excellent Young Scientist Program (grant number: 6119924022, to FW). We would also like to express our gratitude to the participants who took part in the study for their invaluable contributions to this research.

## Author contributions

BW: writing – original draft, formal analysis; YX: writing – review & editing; HW: writing – review & editing, data curation; FW: conceptualization, methodology, writing – original draft, writing – review & editing, funding acquisition.

