## Supplementary Tables 1-3 for "A Critical Rectal Temperature Defines Sex-Specific Leukocyte and Neutrophil Activation During Prolonged Passive Heat Exposure in Young Adults"

**By**

Boan Wei, Yi Xu, Haojian Wang, Faming Wang*

This supplementary file contains 3 tables and 1 figure.

**Supplementary Table S1 |** Sample size, anthropometric characteristics, and baseline physiological parameters for both sexes during six heat exposure scenarios at wet-bulb temperatures ($T_w$) of 32 °C, 33 °C, 34 °C, and 35°C (exposure time: 2.8–8.0 h).

| $T_w$ | $T_w$ = 32°C | | | | | | $T_w$ = 33°C | | | | | |
|---|---|---|---|---|---|---|---|---|---|---|---|---|
| **Conditions** | **$T_{db}$ = 50°C & RH = 24.5%** | | **$T_{db}$ = 40°C & RH = 55.0%** | | **$T_{db}$ = 36°C & RH = 74.5%** | | **$T_{db}$ = 37°C & RH = 74.8%** | | **$T_{db}$ = 42°C & RH = 51.5%** | | **$T_{db}$ = 47°C & RH = 35.5%** | |
| Sex | Males | Females | Males | Females | Males | Females | Males | Females | Males | Females | Males | Females |
| Number | 14 | 10 | 14 | 21 | 10 | 9 | 11 | 11 | 12 | 11 | 12 | 12 |
| | Anthropometric information | | | | | | | | | | | |
| Age (yr) | 24.86 ± 2.32 | 23.00 ± 2.21 | 24.79 ± 1.63 | 23.05 ± 2.11 | 24.60 ± 0.70 | 22.91 ± 1.90 | 24.56 ± 2.30 | 23.44 ± 1.94 | 24.30 ± 2.31 | 23.44 ± 1.94 | 24.25 ± 2.25 | 23.50 ± 1.84 |
| BMI (kg/$m^2$) | 22.01 ± 1.78 | 21.05 ± 2.34 | 21.77 ± 1.33 | 20.10 ± 2.10 | 22.17 ± 1.35 | 20.44 ± 1.84 | 22.17 ± 2.10 | 20.75 ± 2.04 | 22.39 ± 1.50 | 20.75 ± 2.04 | 22.16 ± 1.71 | 20.46 ± 2.12 |
| BSA ($m^2$) | 1.87 ± 0.10 | 1.59 ± 0.10 | 1.90 ± 0.12 | 1.60 ± 0.12 | 1.90 ± 0.14 | 1.59 ± 0.09 | 1.82 ± 0.15 | 1.53 ± 0.11 | 1.83 ± 0.10 | 1.53 ± 0.11 | 1.84 ± 0.10 | 1.52 ± 0.11 |
| | Baseline physiological parameters | | | | | | | | | | | |
| $T_{rec}$ (°C) | 36.87 ± 0.20 | 36.87 ± 0.15 | 36.85 ± 0.22 | 36.95 ± 0.18 | 36.87 ± 0.10 | 6.82 ± 0.16 | 36.86 ± 0.19 | 36.85 ± 0.21 | 36.88 ± 0.19 | 36.88 ± 0.21 | 36.83 ± 0.23 | 36.84 ± 0.17 |
| Leukocytes (×$10^9$×$L^{-1}$)) | 7.57 (5.84, 9.35) | 8.10 (6.88, 10.25) | 8.76 (7.69, 9.50) | 7.59 (6.89, 8.53) | 6.15 (5.33, 7.32) | 6.17 (5.61, 7.31) | 7.67 (6.59, 8.64) | 7.58 (6.52, 8.26) | 7.65 (6.66, 8.69) | 7.67 (6.83, 8.50) | 7.68 (6.73, 8.69) | 7.26 (6.32, 8.18) |
| Neutrophils (×$10^9$×$L^{-1}$) | 3.95 (2.86, 5.43) | 3.86 (3.06, 6.13) | 3.39 (2.69, 4.50) | 3.41 (2.50, 3.76) | 3.01 (2.43, 3.67) | 3.04 (2.48, 3.56) | 3.25 (2.57, 3.53) | 3.52 (2.74, 3.89) | 3.14 (2.85, 4.28) | 3.10 (2.54, 3.78) | 2.69 (2.47, 3.86) | 3.00 (2.74, 3.62) |
| $T_w$ | $T_w$ = 34°C | | | | | | $T_w$ = 35°C | | | | | |
| **Conditions** | **$T_{db}$= 38°C & RH = 75.1%** | | **$T_{db}$ = 43°C & RH = 52.1%** | | **$T_{db}$ = 48°C & RH = 36.2%** | | **$T_{db}$ = 40°C & RH = 70.3%** | | **$T_{db}$ = 45°C & RH = 49.3%** | | **$T_{db}$ = 50ºC & RH = 34.5%** | |
| Sex | Males | Females | Males | Females | Males | Females | Males | Females | Males | Females | Males | Females |
| Number | 11 | 12 | 11 | 11 | 12 | 11 | 14 | 13 | 10 | 13 | 8 | 10 |
| | Anthropometric information | | | | | | | | | | | |
| Age (yr) | 24.25 ± 1.81 | 23.38 ± 1.89 | 24.25 ± 1.81 | 23.36 ± 1.75 | 24.25 ± 1.82 | 23.36 ± 1.75 | 24.85 ± 4.00 | 23.00 ± 2.16 | 24.57 ± 3.98 | 23.00 ± 2.16 | 24.25 ± 1.75 | 22.9 ± 2.47 |
| BMI (kg/$m^2$) | 21.89 ± 1.33 | 21.40 ± 2.19 | 21.89± 1.33 | 21.23 ± 2.21 | 22.66 ± 1.16 | 21.23 ± 2.21 | 22.11 ± 1.77 | 20.91 ± 1.36 | 22.87 ± 0.84 | 20.91 ± 1.36 | 21.56 ± 2.27 | 20.70 ± 1.50 |
| BSA ($m^2$) | 1.83 ± 0.08 | 1.57 ± 0.11 | 1.83 ± 0.08 | 1.57 ± 0.11 | 1.83 ± 0.08 | 1.57 ± 0.11 | 1.85 ± 0.09 | 1.64± 0.08 | 1.86 ± 0.09 | 1.64 ± 0.08 | 1.86 ± 0.10 | 1.63 ± 0.09 |
| | Baseline physiological parameters | | | | | | | | | | | |
| $T_{rec}$ (°C) | 36.78 ± 0.25 | 36.79 ± 0.28 | 36.82 ± 0.22 | 36.95 ± 0.21 | 36.85 ± 0.19 | 36.87 ± 0.25 | 36.86 ± 0.21 | 36.88 ± 0.14 | 36.87 ± 0.20 | 36.81 ± 0.11 | 36.84 ± 0.29 | 36.87 ± 0.26 |
| Leukocytes (×$10^9$×$L^{-1}$)) | 6.56 (6.11, 7.24) | 6.0 (5.55, 7.66) | 6.51 (6.01, 8.37) | 5.29 (4.78,6.69) | 7.42 (6.00, 8.58) | 5.64 (4.92, 7.5) | 5.84 (5.46,8.05) | 6.95 (6.35,7.87) | 6.30 (5.49, 6.88) | 6.57 (4.99,7.59) | 6.80 (5.95,7.15) | 6.78 (6.15, 7.46) |
| Neutrophils (×$10^9$×$L^{-1}$)) | 3.09 (2.88,3.35) | 2.86 (2.15,3.34) | 3.96 (3.34, 4.23) | 2.43 (1.76, 3.57) | 3.73 (2.65, 4.17) | 2.59 (2.43, 3.53) | 2.82 (2.21, 3.50) | 3.19 (2.67,4.14) | 2.80 (1.98,3.53) | 3.11 (2.00,3.44) | 3.06 (2.48, 3.73) | 3.1 (2.72,3.44) |

*Note:* Baseline data for leukocyte and neutrophil data are presented as median [IQR (interquartile range): lower quartile, upper quartile]; all other variables are expressed as mean ± SD.

Abbreviations: $T_w$, wet-bulb temperature; $T_{db}$, dry-bulb temperature; RH, relative humidity; BMI, body mass index; BSA, body surface area; $T_{rec}$, rectal temperature. Twelve conditions were $T_w$-equivalent. For $T_w$ =32–33°C, exposure duration was 8 h; for $T_w$ = 34–35 °C, exposure continued until the rectal temperature reached 38.6 °C.

**Supplementary Table S2 |** Comparison of dehydration rate (%) between sexes.

| $T_w$ (°C) | $T_{db}$ (°C) | RH (%) | Dehydration Rate (%) | |
|---|---|---|---|---|
| | | | Males | females |
| 32 | 50 | 24.5 | 0.54±0.17 | 0.63±0.16 |
| | 40 | 55.0 | -0.30±0.67 | -0.49±0.51 |
| | 36 | 74.5 | 0.35±0.26 | 0.43±0.32 |
| 33 | 37 | 74.8 | 0.27±0.56 | -0.17±0.75 |
| | 42 | 51.5 | 0.76±0.15 | 0.59±0.40 |
| | 47 | 35.5 | 0.55±0.37 | 0.69±0.35 |
| 34 | 38 | 75.1 | -0.52 ± 0.54 | -0.59 ± 0.46 |
| | 43 | 52.1 | -0.06 ± 0.03 | -0.67 ± 0.58 |
| | 48 | 36.2 | -0.05 ± 0.29 | -0.50 ± 0.53 |
| 35 | 40 | 70.3 | -0.47 ± 0.35 | -0.56 ± 1.06 |
| | 45 | 49.3 | -0.60 ± 0.59 | -0.50 ± 0.31 |
| | 50 | 34.5 | -0.25 ± 0.62 | -0.44 ± 0.41 |

The dehydration rate (*DR*) was calculated from Equation (1), expressed as a percentage, which quantifies the degree of dehydration resulting from body water loss during the experiment.

$$DR = \frac{SP + UL - FI}{IBW} \cdot 100\% \quad (1)$$

where, negative DR values indicate a net hydration state (fluid intake exceeds fluid loss), whereas positive values indicate net dehydration. *SP* (sweat production) is the total volume of sweat secreted during the observation period (mL); *UL* (urine loss) is the total volume of urine excreted (mL); *FI* (fluid intake) includes all sources of water uptake (mL); *IBW* (initial body weight) is the participant's body mass before exposure (kg), used to normalize water loss and enable consistent comparisons of relative dehydration.

**Supplementary Table S3 |** Sex differences in baseline leukocyte and neutrophil counts.

| Parameter | Sex | Median (Q1, Q3) ($\times 10^9$/L) | Mann-Whitney U | Z | *r* | *p* |
|---|---|---|---|---|---|---|
| Leukocyte count | Male | 6.76 (5.76, 8.02) | 109518.5 | -1.105 | 0.105 | 0.269 |
| | Female | 6.72 (5.64, 7.86) | | | | |
| Neutrophil count | Male | 3.13 (2.54, 3.76) | 105528.5 | -2.039 | 0.199 | 0.041* |
| | Female | 2.99 (2.43, 3.67) | | | | |

*Note*: Q1, lower quartile; Q3, upper quartile. Units: ×$10^9$/L. *$p < 0.05$ denotes significance, $r$ represents the effect size in the Mann-Whitney U test, and Z represents the standardized test statistic of the Mann-Whitney U test.

### Derivation of the Critical Rectal Temperature ($T_{rec}$) Threshold

The critical rectal temperature ($T_{rec}$) of 38.1 °C identified in this study was determined through a systematic analysis using quadratic polynomial regression models to characterize the relationship between leukocyte increment and $T_{rec}$ in both males and females.

First, separate quadratic polynomial regressions were performed for leukocyte increment versus $T_{rec}$ data in both young males and females to capture the nonlinear relationship. The first-order derivatives of these regression equations were then computed to obtain the instantaneous rate of leukocyte change with respect to $T_{rec}$. Solving for the $T_{rec}$ at which the first-order derivatives for both sexes were equal yielded a crossover temperature of 38.07 °C.

Given the $T_{rec}$ measurement accuracy of ±0.1 °C (YSI 401 rectal probe; see Section 2.2 in the main manuscript) and that temperature data were recorded to one decimal place, 38.07 °C was rounded to 38.1 °C. This temperature represents the threshold differentiating leukocyte response patterns between sexes: below 38.1 °C, females exhibit a higher rate of leukocyte increment, whereas above this threshold, males display a greater and more rapidly accelerating rate. The specific analyses and regression results are presented below.

#### *Quadratic polynomial regression results*

Quadratic polynomial regression of leukocyte increment ($L$) and rectal temperature ($T_{rec}$) data for young males and females yielded the following equations:

$$L_m = a_m \cdot T_{rec}^2 + b_m \cdot T_{rec} + c_m \tag{2}$$

$$L_f = a_f \cdot T_{rec}^2 + b_f \cdot T_{rec} + c_f \tag{3}$$

where, $L_m$ and $L_f$ denote leukocyte increments for males and females ($\times 10^9 \cdot L^{-1}$), respectively; $a_m$, $b_m$, $c_m$ are the regression coefficients for males, and $a_f$, $b_f$, $c_f$ are those for females (exact coefficient values are provided in the Results section of the main manuscript). The quadratic term ($T_{rec}^2$), the linear term ($T_{rec}$), and the constant term together describe the nonlinear relationship between $L$ and $T_{rec}$.

### *Calculation of the first-order derivative function*

To quantify the rate of change of $L$ with respect to $T_{rec}$, the first-order derivatives of Equations (2) and (3) were calculated:

$$\frac{dL_m}{dT_{rec}} = 2a_m \cdot T_{rec} + b_m \tag{1}$$

$$\frac{dL_f}{dT_{rec}} = 2a_f \cdot T_{rec} + b_f \tag{2}$$

The derivative represents the change in leukocyte increment ($L$) per 1 °C change in $T_{rec}$.

### *Calculation of the critical $T_{rec}$*

The critical $T_{rec}$ is defined as the temperature at which the rates of change of leukocyte increment are equal for both sexes, that is, where the first-order derivatives in Equations (4) and (5) are equal:

$$2a_m \cdot T_{rec} + b_m = 2a_f \cdot T_{\mathrm{rec}} + b_f \tag{6}$$

$$T_{rec} = \frac{b_f - b_m}{2a_m - 2a_f} \tag{7}$$

Solving these equations yields a critical $T_{rec}$ of 38.1 °C.

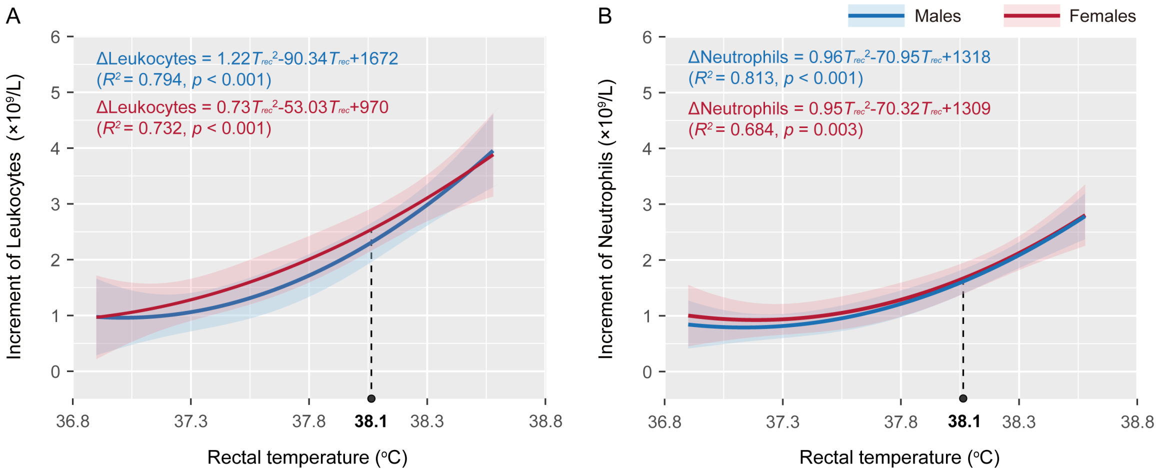


**Supplementary Figure S1** | Critical rectal temperature (38.1°C) as the differentiation threshold for the rate of leukocyte increment between young adult males and females. All data were analyzed using quadratic polynomial regression models.
